# Histone deacetylase inhibitor treatment promotes spontaneous caregiving behavior in C57BL/6J male mice

**DOI:** 10.1101/433995

**Authors:** H.S. Mayer, M. Crepeau, N. Duque-Wilckens, L.Y. Torres, B.C. Trainor, D.S. Stolzenberg

## Abstract

Whereas the majority of mammalian species are uni-parental with the mother solely provisioning care for young conspecifics, fathering behaviors can emerge under certain circumstances. For example, a great deal of individual variation in response to young pups has been reported in multiple inbred strains of laboratory male mice. Further, sexual experience and subsequent cohabitation with a female conspecific can induce caregiving responses in otherwise indifferent, fearful or aggressive males. Thus, a highly conserved parental neural circuit is likely present in both sexes, however the extent to which infants are capable of accessing this circuit may vary. In support of this idea, fearful or indifferent responses toward pups in female mice are linked to greater immediate early gene (IEG) expression in a fear/defensive circuit involving the anterior hypothalamus than in an approach/attraction circuit involving the ventral tegmental area. However, experience with infants, particularly in combination with histone deacetylase inhibitor (HDACi) treatment, can reverse this pattern of neuronal activation and behavior. Thus, HDACi treatment may increase the transcription of primed/poised genes that play a role in the activation and selection of a maternal approach circuit in response to pup stimuli. Here, we asked whether HDACi treatment would impact behavioral response selection and associated IEG expression changes in virgin male mice that are capable of ignoring, attacking or caring for pups. Our results indicate that systemic HDACi treatment induces spontaneous caregiving behavior in non-aggressive male mice and alters the pattern of pup-induced IEG expression across a fear/defensive neural circuit.

## Introduction

*Mus musculus* is a uni-parental rodent species in which the mother solely cares for her young in the wild. However laboratory conditions can contribute to the emergence of maternal-like behavior (pup retrieval, sniffing/licking, crouching) along with the elimination of pup-directed aggression in males (Leblond, 1938; Huck et al., 1982; Jakubowski and Terkel, 1982; Vom Saal and Howard, 1982; Elwood and Ostermeyer, 1984). This is observed in both sexually experienced males, which are often cohabited with females to optimally produce offspring, and sexually naïve male mice, albeit much less frequently. When exposed to pups, sexually naïve male mice tend to show highly variable responses to pups including aggressive, exploratory, avoidant, and even spontaneous caregiving behaviors. Together these data support the idea that the neural circuit that regulates maternal behavior is conserved in male mice, but the extent to which infant stimuli gain access to this circuit varies considerably between individuals due to largely unknown mechanisms.

In females, seminal work uncovering the neural mechanisms that gate infant access to the maternal neural circuit was conducted in postpartum rats (Numan and Stolzenberg, 2008) and recent work has replicated some of these findings in mice (Fang et al., 2018; Kohl et al., 2018). Importantly, motivation to care for offspring first occurs around the time of birth. In non-parental animals, infants activate hypothalamic regions known to regulate anxiety/escape/attack behaviors such as the anterior hypothalamic nucleus (AHN) and ventromedial nucleus of the hypothalamus (VMN) (Sheehan et al., 2000, 2001; Tachikawa et al., 2013). In contrast, the medial preoptic area (MPOA) of the rostral hypothalamus regulates caregiving behavior through its projection to the ventral tegmental area (VTA), which drives the release of dopamine into the nucleus accumbens (NA) causing high levels of maternal responding (Numan and Smith, 1984; Sheehan and Numan, 2002; Numan et al., 2005; Shahrokh et al., 2010). Thus, hormonal stimulation during late pregnancy and birth facilitates the onset of maternal behavior by increasing infant access to this MPOA-VTA-NA circuit. Whereas plasticity within this circuit contributes to the maintenance of caregiving behavior across the postpartum period long after hormonal stimulation has waned (Afonso et al., 2009), caregiving behavior likely depends on changes in both anti-social and pro-social neural systems (Numan and Sheehan, 1997; Kuroda and Numan, 2014). For example, the onset of mothering in rats also coincides with a reduction in the ability of infants to activate fear/defensive neural systems (Sheehan et al., 2000) and experimentally induced reactivation of this system can turn mothering off (Sheehan and Numan, 1997). Thus, the occurrence of caregiving behavior may depend on both a pup-induced activation of the maternal circuit and an inhibition of a competing fear/escape/attack neural system (Numan and Sheehan, 1997).

Whereas the transition from pup avoidance to pup approach is typically linear and uni-directional in female rats, the response of male mice toward pups can be influenced by social or contextual factors. For example, while males transition from aggressive or avoidant responses to approach and caregiving responses following sexual experience (Tachikawa et al., 2013), in the absence of continued pup exposure they can transition back to pup-directed aggression (Vom Saal, 1985). Therefore the male mouse model is useful for understanding the relationship between pup-induced activation of a neural system and pup-directed behavioral responses because males engage in aggressive, avoidant or caregiving responses under predictable circumstances. Recent work supports the idea that the reduced activation of a central aversion system (including AHN/VMN) in response to pups is also associated with the transition to paternal care (Tachikawa et al., 2013; Tsuneoka et al., 2015). Further, expression of the IEG, cFos, within the rhomboid part of the dorsal bed nucleus of the stria terminalis (dBNST) was found to be highly correlated with pup-directed aggression (Tsuneoka et al., 2015), although the mechanism through which activation of a central aversion system mediates distinct types of aversive responses is presently unclear.

Whereas the role of pregnancy hormones in activating the maternal neural circuit has been well described, the mechanisms through which these neural systems are activated to promote caregiving behavior in non-lactating rodents are relatively unknown. Further, how a neural circuit is selected to mediate a specific behavioral response and how factors like sex, experience or reproductive status regulate the selection of a particular circuit over a competing circuit is unclear. One possibility is that transcriptional patterns within specific cell populations program the activation of a particular circuit. Sex may program a particular circuit for default selection from birth.

Reproductive status (sexual experience in males or gestation in females) might re-program the pattern to set a new circuit as default. Repeated experience with pups may lead to neuronal activity-dependent transcriptional changes that result in differential circuit selection (specifically avoidance to approach). Histone deacetylase inhibitor (HDACi) drugs enhance the transcription of genes that are poised or primed for expression (Wang et al., 2009) and in this way may potentiate experience-driven behavioral modifications. Recently, we found that HDACi treatment in virgin female mice increased the likelihood that regions of the maternal neural circuit, rather than regions of the fear/avoidance circuit, were activated during the challenging task of pup retrieval in a novel T-maze (Mayer et al., 2018). Based on this finding, we hypothesized that experience-induced changes in behavioral response selection may depend on the extent to which IEGs are primed within neural regions regulating these responses to pups. Further, HDACi treatment may increase the transcription of primed genes that promote the activation and selection of approach circuits exclusively. Here, we investigate this hypothesis in pup-naïve virgin male mice because of the considerable variation they show in their default behavioral response to pups. To assay region-specific transcriptional response to pups we quantified mRNA expression of two IEGs, *cFos* and neuronal PAS domain protein 4 (*Npas4*) (Sheng and Greenberg, 1990; Sun and Lin, 2016). We measured *Npas4* in addition to *cFos* because unlike *cFos* (Luckman et al., 1994), *Npas4* is exclusively expressed in neurons and is a reliable indicator of neuronal activity (Bepari et al., 2012). In addition, *Npas4* expression has been shown to be critical for plasticity (Ramamoorthi et al., 2011).

## Methods and Materials

### Subjects and drug treatment

All mice were C57BL/6J virgin adult males (45+ days of age) naive to pups, housed on a 12-hour reverse light cycle and given food and water ad libitum. The HDACi sodium butyrate (Sigma-Aldrich) was dissolved in sterile water and was administered at a dose of 8 mg/ml in the drinking water. Control mice received standard drinking water. Drinking water containing sodium butyrate was given ad libitum beginning 24 hours prior to the start of testing and continued throughout testing. Daily drinking water was monitored for all sodium butyrate treated mice. All mice were housed individually for 3-7 days prior to and throughout testing. Behavioral testing was conducted one hour into the dark phase of the light/dark cycle under dim red light. Stimulus pups were obtained from lactating C57BL/6J or CD1 lactating dams in our donor-pup breeding colony. All procedures were in compliance with the University of California, Davis Institutional Animal Care and Use Committee.

### Behavioral Procedures

#### Home cage parental behavior tests

Naïve virgin male mice were treated with sodium butyrate (N= 24) or water (N = 25). Behavioral testing began by scattering three stimulus pups (1-6 days old) in the home cage. Mice were rated using a 5 point scale based on their response to pups during a 15-minute test: 0-repeated biting of pups, 1-rough handled or stepped on pups, 2-spent less than 50% of the test investigating pups, 3-spent more than 50% of the test investigating pups, 4-retrieved at least one pup, 5-displayed full paternal care (retrieval, sniffing/licking and hovering over pups). Mice were then categorized based on their score as aggressive (0-1), indifferent (2), or paternal (4-5). For male mice that were not aggressive toward pups (scores 1-5), latencies to sniff, retrieve each pup to the nest, sniff/lick the grouped pups and hover over pups in the nest were recorded during the 15-minute test. Pups remained in the cage for a total of 2 hours and were then removed and returned to a lactating dam. In the event that a male attacked a pup, the test was stopped and the pups were immediately removed from the cage. Male mice that attacked pups on the first test were not tested again. Males that did not attack pups were tested for 2 consecutive days.

#### Social interaction test

To investigate whether effects of HDACi on behavior are exclusive to interactions with pups, a separate cohort of pup-naïve virgin male mice treated with sodium butyrate (N= 8) or water (N= 8) was tested in the social interaction test. Social interaction testing was conducted in a large Plexiglas open field that contained no bedding (89×63×60 cm), as described previously (Duque-Wilckens et al., 2018). Briefly, the test consisted of 3 consecutive phases (open field, acclimation and interaction). During the open field phase of testing each mouse was introduced into the arena for 3 minutes. Time spent in the center of the arena, corners of the open field and total distance traveled was recorded (Any-Maze, Stoelting). Following the open field phase, a small wire cage was introduced against one wall of the arena (without removing the focal mouse from the arena). During this 3-minute acclimation phase, the time spent within 8 cm of the novel cage (time investigating novel object) or within the 2 corners (8×8 cm each) opposite the wire cage (time away from novel object) was recorded. During the last phase of testing, an unfamiliar same-sex stimulus mouse was placed into the wire cage for 3 min and the time spent investigating the novel mouse was recorded.

### Region-specific Gene Expression in Aggressive and Non-Aggressive Males

Whereas the proportion of animals responding aggressively toward pups was not affected by HDACi treatment, the proportion of animals showing paternal care significantly increased following HDACi treatment. These data suggest that HDACi treatment affects behavioral responses toward pups exclusively in male mice that are not aggressive to pups. In order to distinguish between the effects of HDACi treatment on activity-dependent gene expression in aggressive versus non-aggressive mice, we pre-screened males for their initial behavioral responses toward pups. A single pup was introduced into the cage and behavioral responses were recorded for 15 minutes. Mice that attacked were categorized as aggressive and mice that failed to attack within the 15-minute test were categorized as responsive. In order to match the 30-minute pup exposure time between groups while protecting the pups from infanticide a wire mesh ball (tea infuser; Norpro 1.75 inches in diameter) was used with 50 holes (3mm diameter). Males could make contact with pups but were not able to injure them. All males were habituated to the mesh balls for 48 hours prior to testing. Gene expression was examined in 6 groups: pup-naïve virgin male control mice (N=6), pup-naïve virgin male control mice treated with HDACi (N=7), aggressive virgin males (N=7), aggressive virgin males treated with HDACi (N=7), responsive virgin males (N=9), and responsive virgin males treated with HDACi (N=11). On test day, the ball was removed from the cage and replaced with either a pup or no pup (control).

### Quantification of mRNA by real time PCR

Mice were briefly anesthetized with isoflurane and euthanized by cervical dislocation. Brains were immediately removed, frozen and later sectioned (120 microns) on a cryostat and frost-mounted onto slides. The MPOA (Bregma 0.37 to −0.35), AHN/VMN (Bregma −0.59 to −1.67), and VTA (Bregma −2.69 to −3.51) were dissected out using a blunted 15.5 gauge needle and the dBNST (Bregma 0.49 to −0.35) was dissected out using a blunted 18 gauge needle using coordinates from the Franklin and Paxinos Mouse Brain Atlas. Total RNA was isolated with Qiazol reagent (Qiagen) and purified with an RNeasy^®^ Plus Micro Kit (74004; Qiagen, Valencia, CA) as well as the optional DNase digestion (Qiagen 129046). A Nanodrop™ Spectrophotometer was used to determine the quality (260/280 ratio > 1.8) and quantity of the RNA and poor quality samples were not used. The cDNA templates were prepared using an Applied Biosystems cDNA Synthesis Kit (4368813) according to the manufacturer’s protocol. Quantitative real-time PCR was performed using the ABI Viia7 real-time PCR system. The PCR products of interest were detected using TaqMan^®^ Gene Expression assays from (Applied Biosystems, Carlsbad, CA) (Table 1). All samples were normalized to beta-2 microglobulin (*b2m)*. There were no statistically significant differences in the expression of the endogenous control gene between treatment groups. Target and endogenous control genes were measured in triplicate for each cDNA sample during each real-time run to avoid intra-sample variance. All genes of interest were analyzed with Viia7 Applied Biosystems software using the comparative cycle thresholds (delta delta CT) method. There were no statistically significant differences in relative gene expression between pup-naïve control mice treated with or without sodium butyrate for any gene tested and therefore these groups were collapsed and expression of experimental samples was normalized to the average expression of the combined no-pup control group.

**Table 1.**
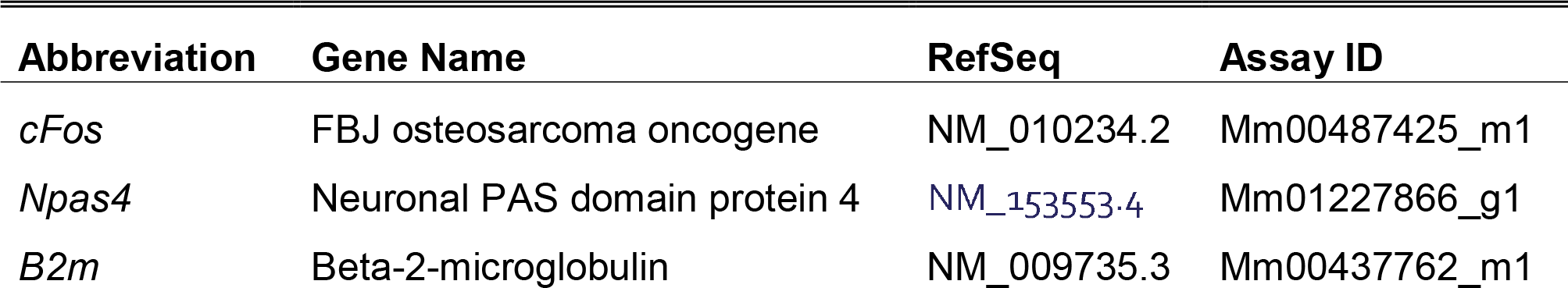
Taqman primers used for qPCR reactions.

| Abbreviation | Gene Name | RefSeq | Assay ID |
| --- | --- | --- | --- |
| *cFos* | FBJ osteosarcoma oncogene | NM_010234.2 | Mm00487425_m1 |
| *Npas4* | Neuronal PAS domain protein 4 | NM_153553.4 | Mm01227866_g1 |
| *B2m* | Beta-2-microglobulin | NM_009735.3 | Mm00437762_m1 |

**Table 2.**
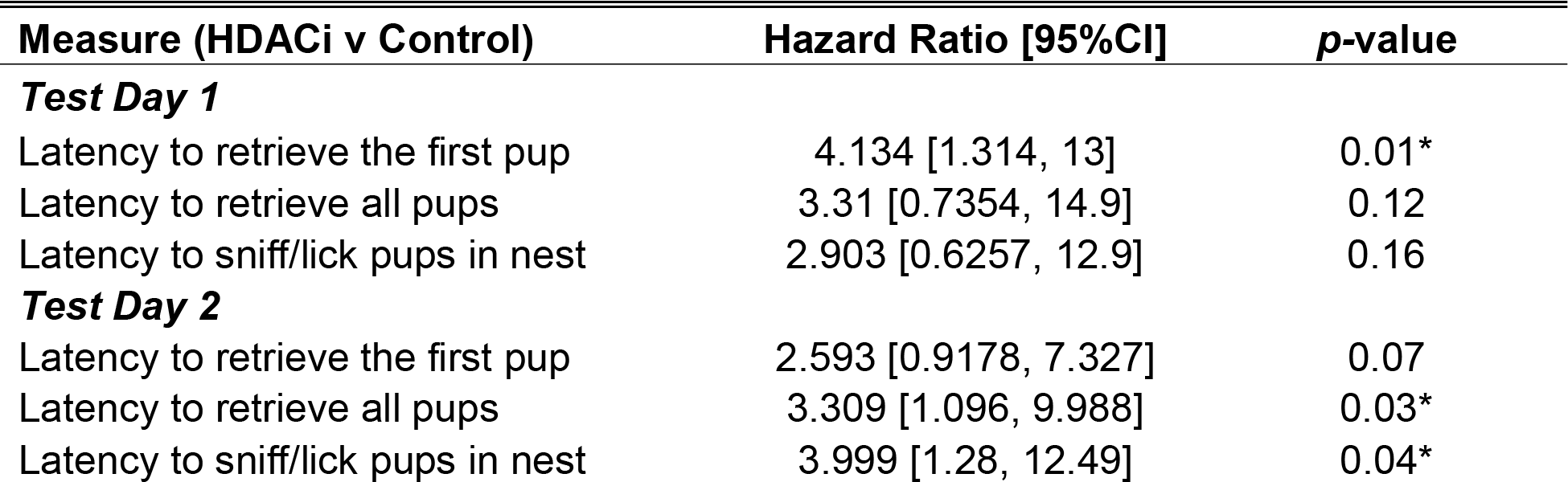
Hazard ratios, calculated from all the data in the survival curves for pup retrieval and pup licking behaviors, indicate the rate at which one group retrieves or licks pups compared to the other.

### Serum Testosterone Assay

To assess whether HDACi treatment could have affected circulating levels of testosterone at the time of pup presentation, a separate cohort of pup-naïve virgin male mice was treated with sodium butyrate (N=7) or water (N=7) for 24 hours. Cardiac blood was collected at the time when pups would have been presented (1 hour after lights go off). Blood was left to coagulate at room temperature for ≥ 30 minutes before centrifugation at 3000g for 10 minutes at 4 degrees Celsius. Supernatant was transferred to a clean microcentrifuge tube and stored at −80 degrees Celsius until assayed. A DRG ELISA kit (EIA-1559) was used to assay serum testosterone according to the manufacturer’s protocol. The manufacturer reports the monoclonal antibody has a dynamic range between 0.083 and 16 ng/mL and the intra assay variance across an n of 20 is 4.16%, 3.28%, and 3.34% at low, mid, and high concentrations, respectively. A standard curve was fit using the 4-parameter logistics method. Experimental samples were assayed in triplicate on a single plate and the intra assay variance was 3.41%.

### Statistical Analysis

Probability data were analyzed using Chi Square and Fisher’s Exact tests. The frequency of pup retrieval (number of pups retrieved) was analyzed by a mixed two-way ANOVA (Treatment x Time), with repeated measures on the second factor. Latency to the first pup contact (sniff) on the first test day was analyzed using a student’s T-test because all of the subjects completed the task in the duration of the test. Survival analyses were used to analyze all other latency data (pup retrieval and sniff/lick) (Jahn-Eimermacher et al., 2011). This method takes into account that some subjects did not retrieve pups during the 15-minute test and censor those data. These latency data are plotted using Kaplan–Meier survival curves in which the fraction of mice that have retrieved (or sniff/licked) pups at each time point is calculated using the product limit (Kaplan-Meier) method. We used the Mantel-Cox log-rank test to statistically compare survival curves on each test day. In addition, hazard ratio and confidence intervals are reported for each variable. The hazard ratio, which is calculated from all the data in the survival curve, indicates the rate at which one group retrieves or licks pups compared to the other. Relative gene expression data were analyzed using two way ANOVAs (Behavior X Treatment). To determine whether IEGs were induced relative to no-pup controls, a one-sample T-test was used to compare each group to the hypothetical value “1”. All other experiments comparing two independent groups were analyzed using a student’s T-test. All statistical tests were two tailed. For ANOVA data, planned comparisons were analyzed using Fisher’s LSD post hoc tests. All data were analyzed using GraphPad Prism 7 software (GraphPad, Inc., La Jolla, CA).

## Results

### Effects of HDACi on Behavioral Response to Pups

HDACi treatment significantly affected the probability of aggressive, indifferent or paternal responses toward pups in virgin male mice [Χ^2^ (2) = 7.906, p = 0.0192, V=0.40; Fig. 1]. Specifically, HDACi treatment induced spontaneous paternal behavior in non-aggressive male mice (indifferent versus paternal, p=0.0108, Fisher’s Exact Test). All non-aggressive males retrieved more pups as a result of pup experience [main effect of time, F(1,17) = 6.434, p = 0.0213, η^2^ = 0.12] and HDACi treated males retrieved more pups than control males [main effect of treatment F(1,17) = 10.95, p = 0.0042, η^2^ = 0.22], particularly on the first test day (p<0.05, d = 1.22). There were no significant differences in latency to first approach pups on test day 1. HDACi treated males were faster to retrieve the first pup on test day 1 [Χ^2^ (1) = 5.894, p = 0.0152, HR 4.134; 95% CI,1.314, 13.00; Fig. 2]. On the second test day, HDACi treated males were faster to retrieve all pups to the nest [Χ^2^ (1) = 4.506, p = 0.0338, HR 3.309; 95% CI, 1.096, 9.988] and lick pups in the nest [Χ^2^ (1) = 5.689, p = 0.0171, HR 3.999; 95% CI, 1.280, 12.49] when compared to control males.

**Figure 1.**
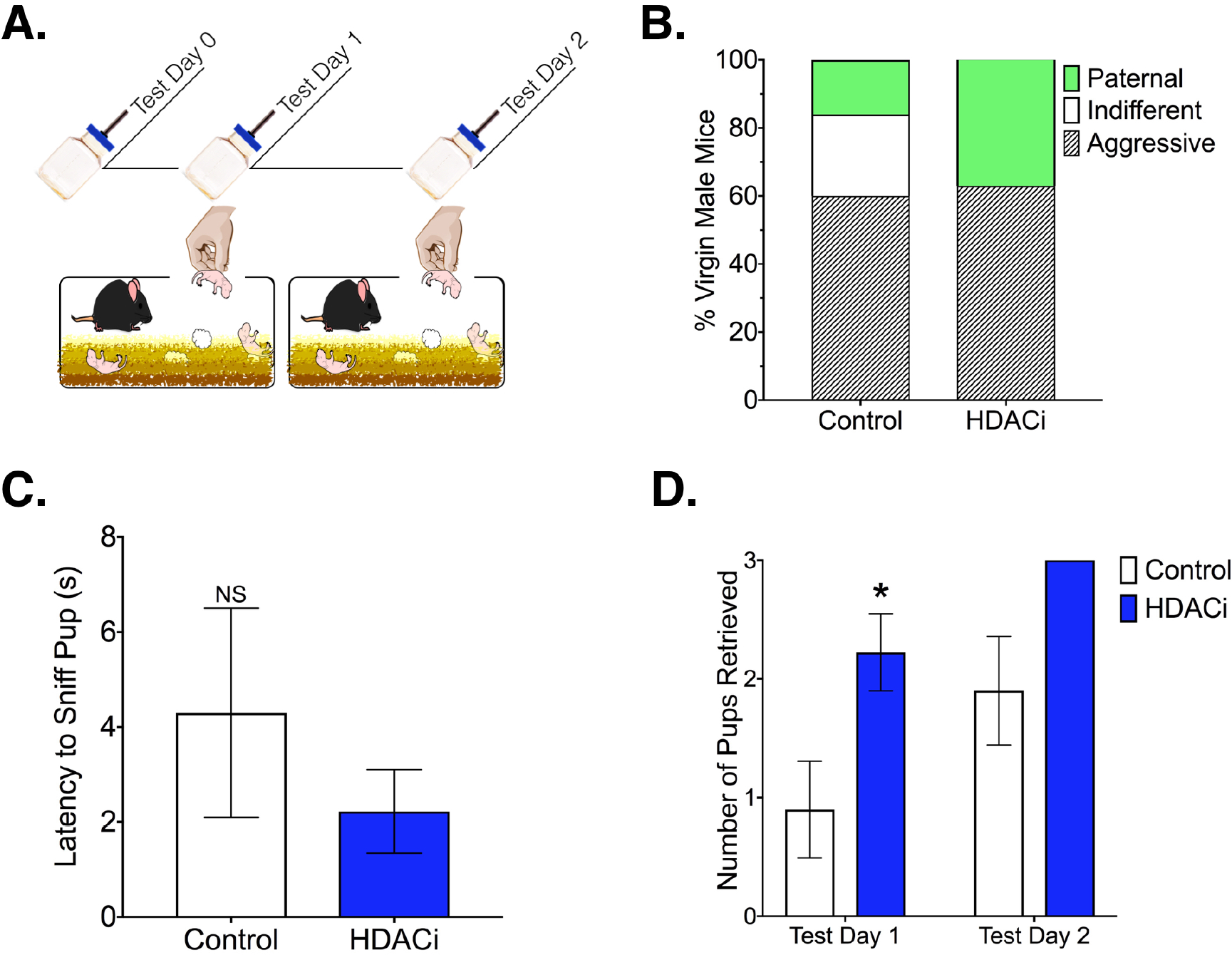
Effects of HDACi treatment on behavioral response selection in the home cage. **A.)** Timeline for Experiment 1: Mice were treated with HDACi or water for 24 hours prior to the start of testing. Only mice that did not show pup-directed aggression were tested on day 2. **B.)** Probability of behavioral response to pups varied significantly by treatment (Fisher’s Exact Test, p = 0.02). All non-aggressive HDACi treated mice showed spontaneous caregiving behavior compared to 40% of control mice (Fisher’s Exact Test, p = 0.01). **C.)** Mean ± SE latency to approach and contact a pup on the first test day did not vary by HDACi treatment. **D.)** HDACi treated males retrieved more pups than controls and all males showed experience-induced improvements in retrieval (Main effects of treatment and time). *Significantly different than control, planned comparison, p< 0.05, d = 1.22

**Figure 2.**
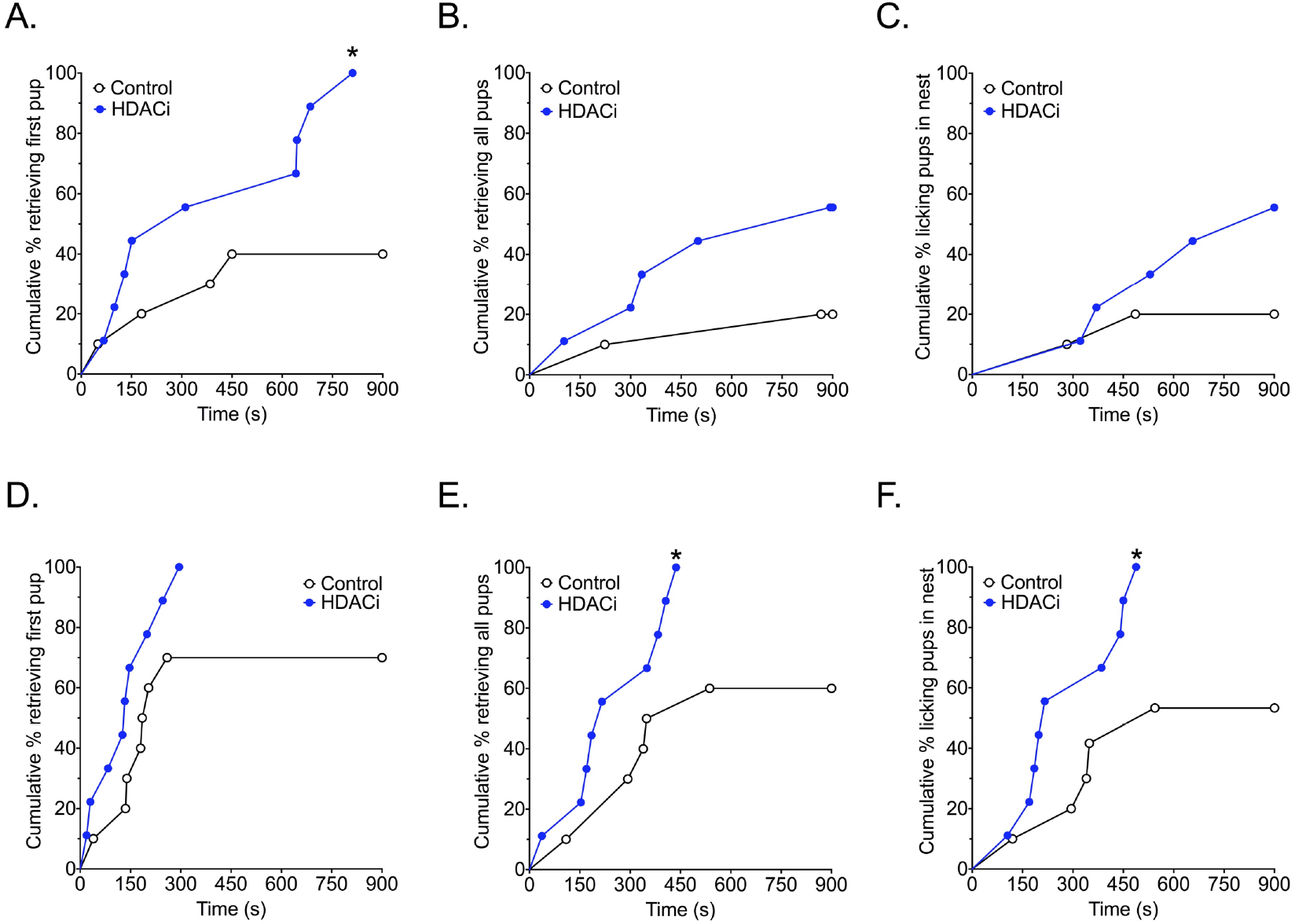
Effects of HDACi treatment in non-aggressive males on the latency to respond to pups. Kaplan-Meier survival curves show the proportion of animals completing the retrieval tasks (retrieving first or last pup) or licking retrieved pups in the nest at each time point on the X-axis in the home cage. **A-C)** HDACi treated male mice were faster to retrieve the first pup on test day 1. **D-E)** HDACi treated males were faster to retrieve all pups and lick retrieved pups in the nest on test day 2. *Significantly different from control group, Chi Square tests, p< 0.05.

### Effects of HDACi on Social Interaction with a Novel Adult Conspecific

There were no significant differences in locomotion (total distance travelled, p = 0.964), thigmotaxis (time in the corners, p = 0.5025) or exploration (time in the center, p = 0.5256) during the open field phase of the social interaction test (Fig. 3). During the acclimation phase, HDACi treated mice spent more time in the corners [t(14)= 2.307, p = 0.0369, d = 1.23] and less time investigating the novel empty cage [t(14)= 2.2448, p = 0.0282, d = −1.31]. However, during the social interaction phase of the test, there were no group differences in locomotion (p = 0.3777), thigmotaxis (time in corners, p = 0.4177) or social interaction time (p = 0.6552).

**Figure 3.**
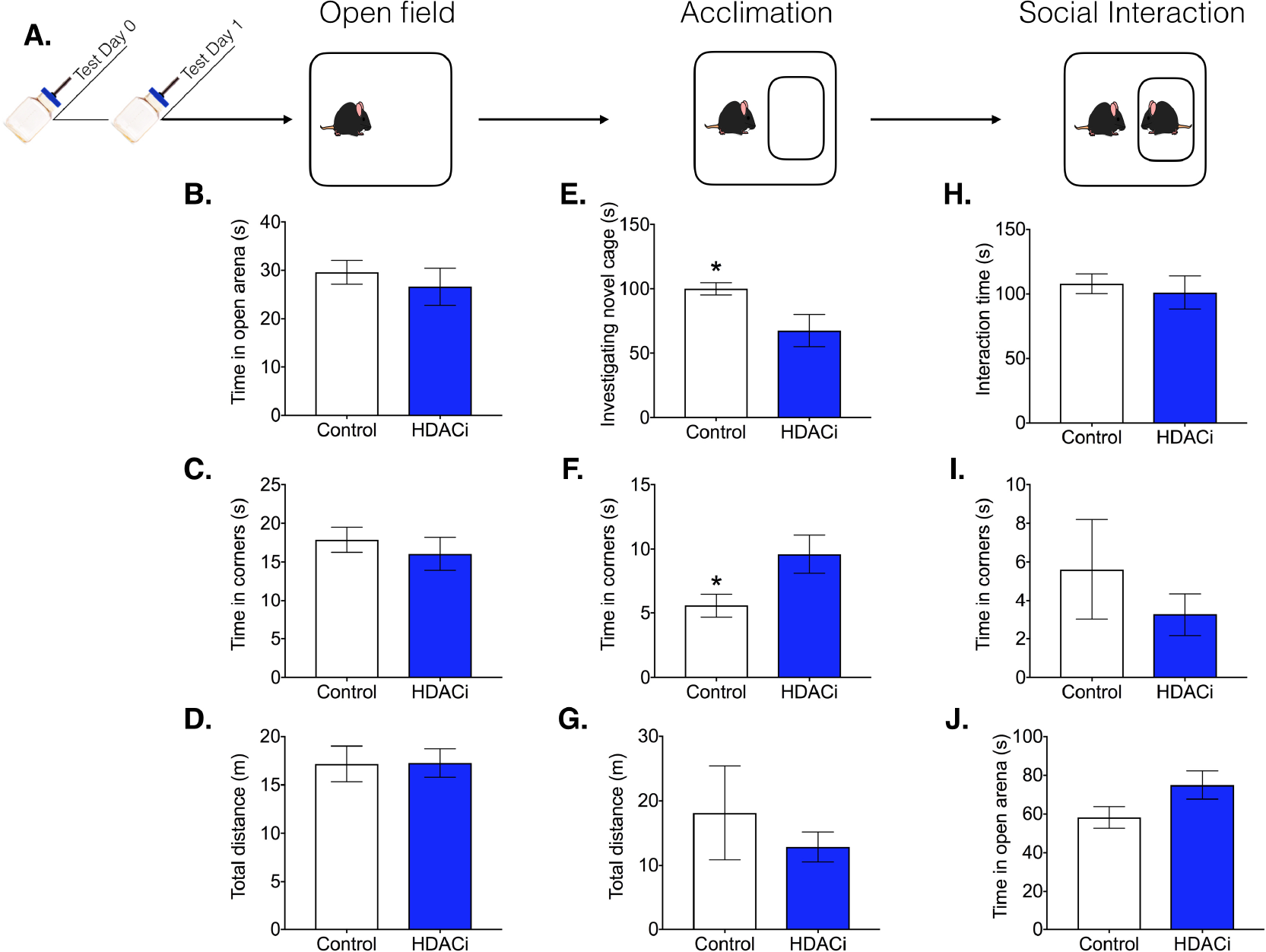
HDACi treatment had no effect on social behavior. **A.)** Timeline for Experiment 2: Males were treated with given sodium butyrate in the drinking water 24 hours before the start of testing. The social interaction test consisted of 3 phases (each lasting 3 min). All data are presented as Mean± SE. **B-D)** Activity during the 3 min open field test was not altered by HDACi treatment. **E-G)** Upon introduction of a novel empty cage, HDACi treated males spent significantly more time in the corners of the arena and less time investigating the empty cage. **H-J)** HDACi treatment had no effect on activity or investigation of a novel adult conspecific *Significantly different from control group, p< 0.05, d_s_ > 1.2.

### Effects of HDACi on Circulating Testosterone

We tested the possibility that effects of HDACi treatment on spontaneous caregiving behavior were related to a treatment-induced change in the circulating level of testosterone by assaying plasma testosterone in male mice exposed to sodium butyrate (or regular water) for 72 hours (Fig. 4). There was no significant effect of HDACi treatment on testosterone levels in virgin male mice (p = 0.2728)

**Figure 4.**
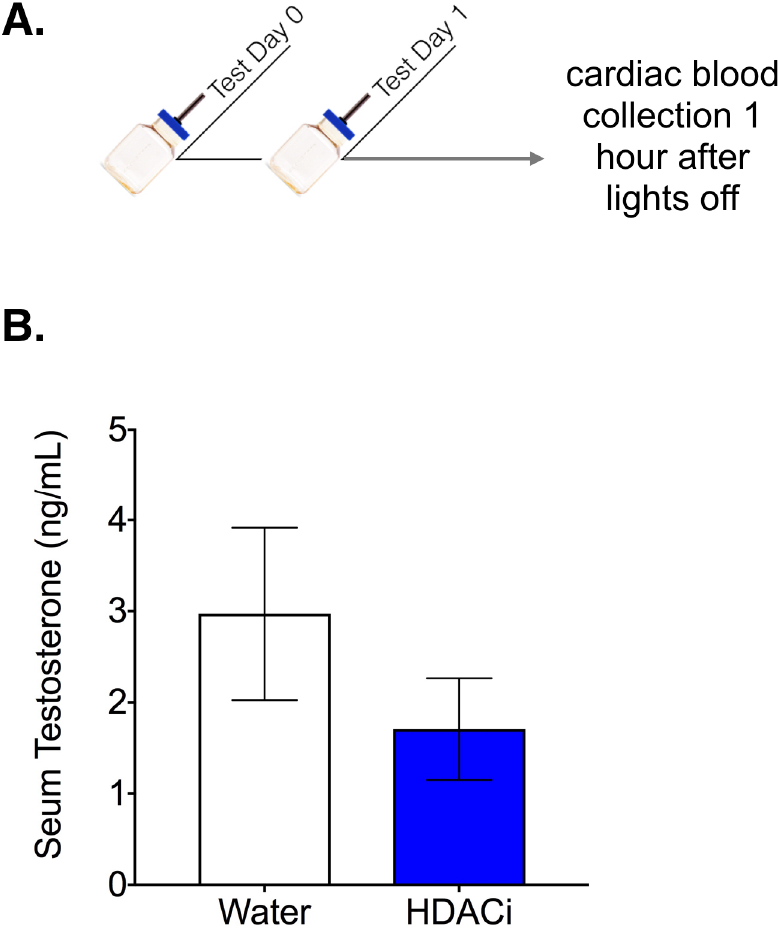
HDACi treatment had no effect on serum testosterone. **A.)** Timeline for Experiment 3: Males were given sodium butyrate in the drinking water 24 hours before cardiac blood collection. **B.)** Concentration of testosterone was not significantly different between groups (p = 0.27)

### Effects of HDACi on Activity-Dependent Gene Expression in Approach/Avoidance Nodes

#### cFos mRNA expression

*cFos* expression was significantly induced by pup exposure in all male mice within the MPOA, AHN/VMN and dBNST regardless of behavioral group or treatment (one sample T-test for each condition in each region, p< 0.05, d_s_ > 0.5; Fig. 5). *cFos* expression in the VTA was significantly higher in aggressive versus responsive males, regardless of HDACi treatment [Main effect of behavioral predisposition: F(1,24) = 4.762, p = 0.039, η2 = 0.16]. In fact, pup-induced *cFos* expression failed to reach statistical significance in responsive males compared to an empty mesh ball (p = 0.07, d = 0.75). In the AHN/VMN, there was a significant interaction effect between behavioral predisposition and HDACi treatment in relative *cFos* expression [F(1,24) = 6.714, p = 0.016, η2 = 0.22]. HDACi treatment reduced *cFos* expression in the AHN/VMN in males that were responsive, but not aggressive, toward pups (p< 0.05, d = −1.10).

**Figure 5.**
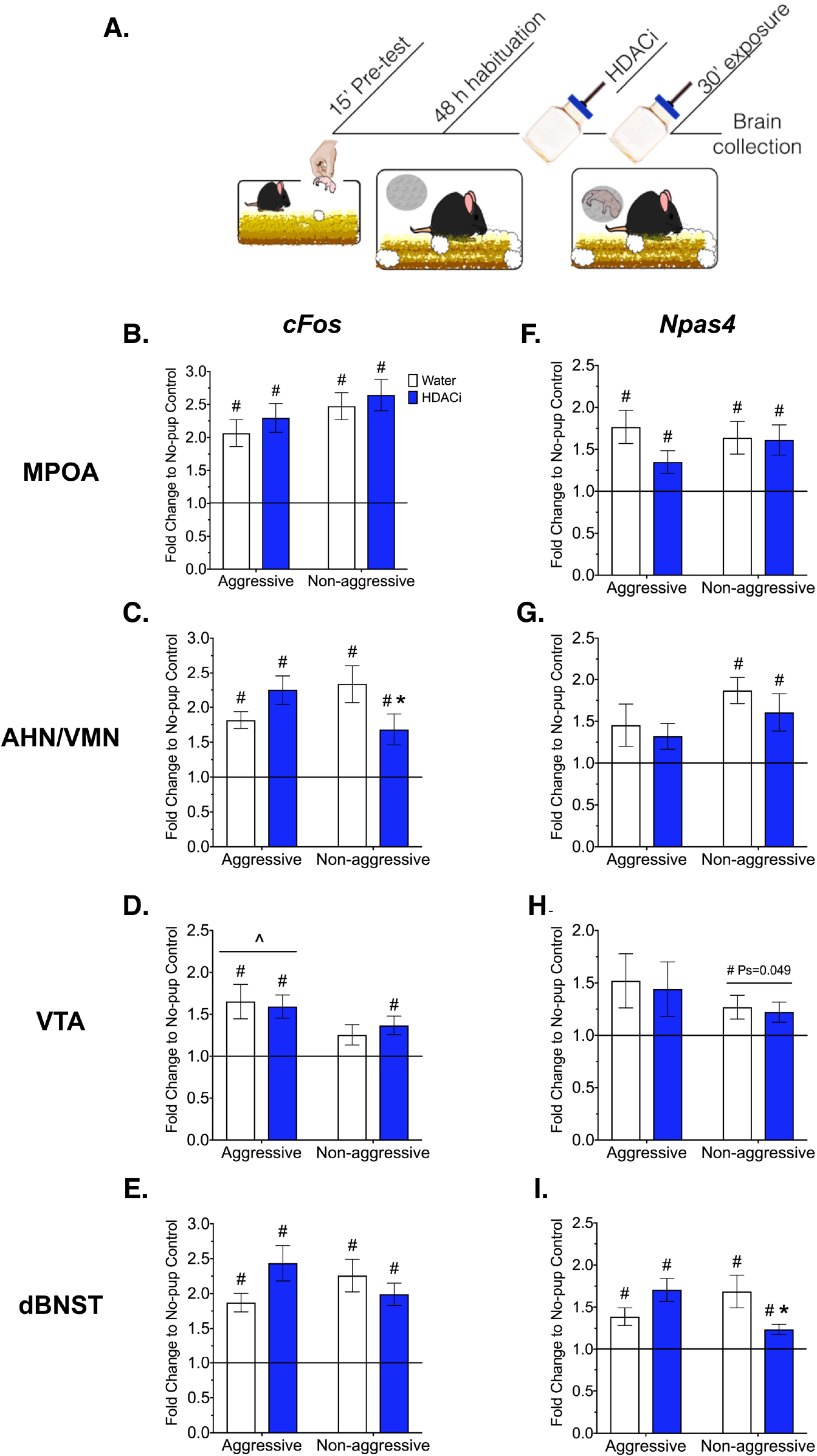
Effects of HDACi treatment on IEG expression. **A.)** Timeline for Experiment 4: Males were given a brief pre-test to identify aggressive or responsive behaviors toward pups. Following the pre-test, males were habituated to the mesh ball for 48 hours. Twenty four hours prior to pup exposure, males were given HDACi-treated or regular water. On test day, a pup was placed into the mesh ball for 30 min before brain collection. Mean± SE *cFos* **(B-E)** and Npas4 **(F-I)** mRNA expression within neural regions associated with pup avoidance/approach. The black line represents normalization to the no-pup control group. #Significantly different from no-pup control, One sample T-tests, p< 0.05 ^Main effect of behavior, 2 way ANOVA, p< 0.05 *Significantly different from corresponding control-treated group, Planned Comparison, p< 0.05.

#### *Npas4* mRNA expression

The immediate early gene, *Npas4*, was also significantly induced by pup exposure in all male mice regardless of behavior or HDACi treatment, but only within the MPOA and dBNST, (one sample T-test for each condition in each region, p< 0.05, d> 0.9). Within the VTA, *Npas4* induction was related to behavioral predisposition, with only responsive mice showing a significant elevation of *Npas4* over no-pup control (p< 0.05, d> 0.7). Similarly, induction of *Npas4* by pup-exposure was limited to responsive mice within the AHN/VMN (p < 0.05,d > 0.9). Within the dBNST, behavioral predisposition and HDACi treatment interacted to affect *Npas4* expression [F(1,29) = 7.569, p = 0.01, η^2^ = 0.28]. HDACi treatment significantly reduced *Npas4* expression in responsive males (p< 0.05, d = −1.13).

## Discussion

Five important conclusions emerge from the results of the present study. First, HDACi treatment induced spontaneous caregiving behavior over pup avoidance, but has no effect on pup-directed aggression. For non-aggressive males, HDACi treatment reduced the latency to retrieve the first pup and increased the number of pups retrieved within 15 minutes of the first pup exposure. In addition to its effects on spontaneous care, HDACi treatment also amplified experience-induced changes in caregiving behavior. HDACi treated males were faster to group pups and lick pups in the nest compared to non-aggressive controls on test day 2. Second, the prosocial effects of HDACi treatment may be specific to pups because HDACi did not affect social investigation of an adult male conspecific. Further, HDACi treatment did not produce a reduction in general fearfulness as measured by exploration of a novel environment. If anything, HDACi treatment was associated with an avoidance of novel objects. Third, the induction of spontaneous caregiving behavior by HDACi treatment was probably not related to a reduction in testosterone because HDACi treatment had no effect on circulating levels of testosterone. Fourth, in line with the finding that HDACi treatment produces behavioral effects exclusively in non-aggressive mice, the effects of HDACi treatment on IEG expression were also limited to non-aggressive males. For example, *cFos* expression in response to pup cues was reduced in HDACi-treated non-aggressive males within the AHN/VMN. Further, HDACi treatment significantly reduced *Npas4* expression in the dBNST, a region that includes the rhomboid nucleus, which may interfere with caregiving behavior through its direct inhibition of the central MPOA (Tsuneoka et al., 2015). In contrast, no effects of HDACi treatment on IEG expression were detected within neural regions associated with pup approach. Both *cFos* and *Npas4* were uniformly induced in the MPOA in all mice exposed to pups. In the VTA, *cFos* expression was induced in mice that show motivated behavioral responses toward pups (regardless of whether that response was pro or anti-social) and surprisingly *cFos* was higher in males that showed pup-directed aggression. *Npas4* induction in the VTA, on the other hand, was limited to non-aggressive males. Finally, the two IEGs examined, *Npas4* and *cFos*, did not show the same pattern of expressions in response to the same pup cues in most of the regions examined. Thus, further investigation of *Npas4* in response to pup cues within these circuits may provide new insight into mechanisms of parental care and experience-induced plasticity.

The behavioral results presented here for control-treated male mice are consistent with other reports of the highly variable response of virgin C57BL/6J mice to foster pups. The fact that the facilitatory effects of HDACi treatment on parental behavior were limited to non-aggressive mice suggests that there is an interaction between individual variation in response to pups and HDACi treatment. Although there is some evidence for a developmentally regulated onset of aggression in C57BL/6J mice, factors that contribute to the individual variation in the response of sexually naïve adult male mice to pups are mostly unknown (Svare and Mann, 1981; Amano et al., 2017). In general there is weak support for a relationship between circulating testosterone and paternal responsiveness in rodents (Wynne-Edwards and Timonin, 2007), although castration does reduce infanticide in virgin male mice (Gandelman and Vom Saal, 1975). However, in the present study we found no evidence for effects of HDACi treatment on circulating levels of testosterone. These data fit with the finding that HDACi treatment also had no effect on aggressive behavior in virgin male mice.

The finding that HDACi treatment promotes caregiving behavior exclusively in non-aggressive males is consistent with our previous work, which has reported the facilitatory effects of HDACi in virgin female mice, which are typically non-aggressive (Stolzenberg et al., 2012, 2014; Mayer et al., 2018). Together these findings suggest that HDACi treatment acts on a conserved neural substrate. However, the extent to which HDACi treatment would fail to promote caregiving behavior in aggressive female mice is unknown. It should be noted that when rare instances of infanticide have occurred we have not found differences between HDACi treated and control female mice (unpublished findings). The effects of HDACi treatment on caregiving behavior exclusively in non-aggressive mice suggest that the molecular pathways, which might be responsible for promoting parental care, are accessible and perhaps primed or poised in non-aggressive males. This idea is related to the known action of HDACi on the expression of transcriptionally poised genes due to the recruitment of HDACs to transcriptionally active sites across the genome, implying that HDACs act as a braking mechanism on activity-induced gene expression rather than a mechanism for gene silencing (Wang et al., 2009).

In the present study we examined the expression of *cFos* and *Npas4*, two IEGs that show stimulus-driven transient expression in the brain. Although *Npas4* expression has never been examined in response to pup cues, *cFos* expression (both mRNA and protein) has been investigated extensively in response to pup cues in both male and female mice as well as female rats (Numan and Numan, 1997; Numan et al., 1998; Sheehan et al., 2000; Tachikawa et al., 2013; Tsuneoka et al., 2013, 2015; Kohl et al., 2018). Our data indicate a near global induction of *cFos* in response to pup cues, and this finding is consistent with other reports that have identified the MPOA, AHN, VMN, and dBNST as regions that are sensitive to pup stimulation. However, the finding that *cFos* induction in these regions was not, for the most part, related to the behavioral response to pups (as determined in the pre-test) was quite surprising. For example, we hypothesized that *cFos* in the AHN/VMN and dBNST would be exclusively induced in aggressive male mice, whereas *cFos* induction in the VTA and MPOA would be limited to non-aggressive males. Further, we predicted that HDACi treatment would amplify the *cFos* response in the MPOA and VTA of pup-responsive males. These hypotheses were based on previous reports of differential Fos expression in sexually naïve (aggressive) males compared with sexually experienced (paternal) C57BL/6J males.

For example, compared to sexually experienced males, aggressive virgins had an exclusive induction of cFos in cells of the AHN, ventrolateral VMN, as well as some subregions of dBSNT (Tachikawa et al., 2013), and although Fos was induced (relative to non-pup control) in paternal males within some subregions of the dBNST, the Fos response of aggressive males was higher. In contrast, we did not find an exclusive relationship between *cFos* induction in the AHN/VMN and aggressive behavior. Instead, *cFos* was induced relative to no-pup control in all male mice exposed to pups. However, it should be noted that our study assayed sexually naïve males with spontaneous aggressive and non-aggressive responses to pups, whereas Tachikawa et al (2013) examined paternal males that had experience caring for pups prior to examining pup-induced Fos response. Certainly virgin males showing spontaneous care are less responsive to pups than pup-experienced fathers. Thus, one possibility is that as caregiving behavior increases the ability of pups to induce a Fos response in the AHN/VMN decreases. In support of this idea, males in our study that would have been most responsive to pups (those treated with an HDACi) showed significantly less *cFos* expression in the AHN/VMN in response to pup cues. Although it is unclear whether males without mating or pup experience would have also failed to show a pup-induced Fos response in the AHN/VMN, a recent investigation examined pup-induced Fos expression within multiple subregions of the dBNST and MPOA in sexually naïve male mice that were aggressive or spontaneously parental (Tsuneoka et al., 2015). This work reported that the number of Fos positive cells in the central part of the MPOA and the rhomboid nucleus of the dBNST were highly predictive of paternal or infanticidal responses, respectively. However, when pup cues were presented indirectly (pups presented in a mesh ball) the differences in Fos expression between parental and infanticidal males in all subregions of the MPOA were eliminated and only the rhomboid and anterior lateral parts of the BNST were found to be significantly different between these groups. Our dBNST tissue punches included several subregions of dBNST in addition to the rhomboid/anterior lateral subregions and therefore any effect of the rhomboid region alone may have been washed out.

To our knowledge, this study is the first to examine differential *cFos* expression in aggressive and responsive virgin males within the VTA. Our hypothesis that non-aggressive males would have higher pup-induced *cFos* expression in the VTA was based on data from virgin female mice (Mayer et al., 2018). Again we were surprised to find that *cFos* was induced in both aggressive and non-aggressive HDACi-treated males. Further, aggressive males had significantly higher expression than non-aggressive males, regardless of HDACi treatment. One interpretation is mice that are least likely to approach and interact with pup (non-aggressive control-treated males) do not show a *cFos* response to pup cues in the VTA. Thus, motivation to approach pups, regardless of the intent to kill or care, is associated with *cFos* induction. In support of this idea, optogenetic stimulation of MPOA neurons that project to the VTA increased motivation to reach pups (by climbing over a physical barrier) in male and female mice even though males killed pups once they came into contact with them (Kohl et al., 2018). Therefore, perhaps it’s not surprising that *cFos* expression alone in the VTA doesn’t predict the intention to kill or care for pups. Together these findings fit nicely with the idea that hypothalamic interaction with the mesolimbic dopamine system regulates social motivation more broadly, including approach responses toward both appetitive and aversive (Numan, 2014). It should be noted that HDACi-treated non-aggressive males have significantly reduced *cFos* expression in the AHN/VMN coupled with a pup-induced *cFos* response in the VTA, whereas non-aggressive control-treated males have a significantly higher *cFos* response in the AHN but no pup-induced *cFos* response in the VTA. Thus, perhaps it is the combination of these responses that is important for caregiving behavior.

In addition to *cFos*, we chose to examine *Npas4*, another IEG with a similar time course of induction to *cFos* (Ramamoorthi et al., 2011). Whereas *cFos* transcription is induced in brain cells by a number of different extracellular stimuli, *Npas4* induction is specifically linked to depolarization of neurons, and therefore may provide some indication of the neuronal response to pups within these regions (Lin et al., 2008). Further, *Npas4* expression is induced in response to learning, rather than exposure to novel or robust stimuli. For example, *Npas4* is induced in the hippocampus following contextual fear learning but unlike *cFos*, *Npas4* expression is not induced by shock alone (Ramamoorthi et al., 2011). Once translated, Npas4 protein serves as a transcription factor, regulating the expression of several late-responding genes that are also critical for neuronal plasticity and particularly new synapse formation (Sun and Lin, 2016). Thus, stimulus-induced expression of *Npas4* might suggest that a neuronal response to pups rather than an increased input to cells as a result of pup exposure.

Our data indicate that *Npas4* induction was limited to non-aggressive males in both the AHN/VMN and VTA, although the VTA data may be interpreted with some caution as this result barely reached statistical significance. HDACi treatment was without effect on *Npas4* expression in these sites, thus *Npas4* induction in these regions might be linked to the non-aggressive behavioral response rather than caregiving behavior per se. With respect to HDACi-induced changes in *Npas4* expression, the dBNST was the only site affected. Therefore the HDACi-induced reduction in *Npas4* expression may be related to the induction of paternal care. Finally, the dBNST and the MPOA may be particularly sensitive to pup stimuli given that we found a significant induction of both *Npas4* and *cFos* in all males exposed to pups. The fact that HDACi treatment significantly lowered *Npas4* in the dBNST fits with the idea that this region plays an inhibitory role in parental behavior, although the present data are not consistent with the idea that this role involves the exclusive regulation of pup-directed aggression.

In conclusion, the results of the present study indicate that HDACi treatment can induce spontaneous caregiving behavior in non-aggressive male mice. The facilitatory effect of HDACi treatment is robust and specific to parental behavior. All non-aggressive males with HDACi treatment responded to pups within 15 minutes of pup exposure HDACi treatment did not reduce neophobia or increase social behavior generally. HDACi-induced reduction in IEG expression within two sites that inhibit caregiving behavior is likely related to the induction of spontaneous caregiving behavior, however the overall pattern of pup-induce IEG expression was not entirely supported by our predictions. An aggressive behavioral predisposition was not associated with the exclusive expression of *cFos* in regions of the brain linked to fearful/defensive behavior in response to pup cues. Similarly, we did not find greater activation of IEG expression in the MPOA of non-aggressive males in response to pup cues. Together these findings underscore the importance of understanding how the MPOA and its interaction with downstream neural sites regulate spontaneous care, indifference or pup-directed aggression. Finally, the present data support the idea that *Npas4* expression may be a more specific marker for neuronal activation, as unlike cFos expression, *Npas4* was differentially expressed in non-aggressive and aggressive mice. Future work will need to gain a cellular resolution of *Npas4* activity in these regions in order to better understand its role in paternal experience-induced plasticity.

## Acknowledgements

This work was supported by the National Institute of Child Health and Human Development [1R01HD087709-01A1] and the National Institute of Mental Health [R01MH103322].

## Notes

**Disclosure of Conflicts of Interest:** The authors declare no conflicts of interest.

